# Multi-ATOM: Ultrahigh-throughput single-cell quantitative phase imaging with subcellular resolution

**DOI:** 10.1101/510693

**Authors:** Kelvin C. M. Lee, Andy K. S. Lau, Anson H. L. Tang, Maolin Wang, Aaron T. Y. Mok, Bob M. F. Chung, Wenwei Yan, Ho Cheung Shum, Kathryn S. E. Cheah, Godfrey C. F. Chan, Hayden K. H. So, Kenneth K. Y. Wong, Kevin K. Tsia

## Abstract

A growing body of evidence has substantiated the significance of quantitative phase imaging (QPI) in enabling cost-effective and label-free cellular assay, which provides useful insights into understanding biophysical properties of cells and their roles in cellular functions. However, available QPI modalities are limited by the loss of imaging resolution at high throughput and thus run short of sufficient statistical power at the single cell precision to define cell identities in a large and heterogeneous population of cells – hindering their utility in mainstream biomedicine and biology. Here we present a new QPI modality, coined multi-ATOM that captures and processes quantitative label-free single-cell images at ultra-high throughput without compromising sub-cellular resolution. We show that multi-ATOM, based upon ultrafast phase-gradient encoding, outperforms state-of-the-art QPI in permitting robust phase retrieval at a QPI throughput of >10,000 cell/sec, bypassing the need for interferometry which inevitably compromises QPI quality under ultrafast operation. We employ multi-ATOM for large-scale, label-free, multi-variate, cell-type classification (e.g. breast cancer sub-types, and leukemic cells versus peripheral blood mononuclear cells) at high accuracy (>94%). Our results suggest that multi-ATOM could empower new strategies in large-scale biophysical single-cell analysis with applications in biology and enriching disease diagnostics.

## 1. Introduction

Quantitative phase imaging (QPI) allows label-free, non-invasive quantification of local optical path length of biological specimens [1–3] that can be harnessed to gain insight into the cellular biophysical properties. This capability has advanced our understanding in cell biology and has enabled novel strategies in biomedical diagnostics. For examples, cell size and mass evaluated from the quantitative phase are linked to cellular progression and apoptosis [4, 5]. Subtle changes in sub-cellular organizations visualized in QPI can be used as a hallmark of cancer [6–8]. When it comes to probe the biophysical properties at single-cell precision, QPI could thus be of functional significance in complementing the current state-of-the-art single-cell analysis (SCA) [9, 10] in a cost-effective manner.

Unfortunately, widespread utility of QPI for biophysical phenotyping in SCA remains limited. The major challenge is that current QPI methodologies lack the ability to simultaneously deliver the two critical, but often conflicting attributes, i.e. *sub-cellular resolution* and *high imaging throughput*, both of which are essential for assessing cell population heterogeneity with high discriminatory power. In the current QPI approaches, the sub-cellular resolution is commonly achieved at the expense of the imaging throughput which is limited to only few tens to hundreds of cells within a field-of-view (FOV) [11]. Despite that the throughput problem can partially be alleviated through automated sample scanning (up to ~10^4^ cells) [12], it is further exacerbated by the inherent limitations of current camera technology, i.e. severe image motion blur, degradation in image resolution and loss of detection sensitivity at high speed. This also explains the limited flow rate, and thus single-cell imaging throughput in the available QPI flow cytometers [13–15]. Even if they were integrated with the state-of-the-art imaging flow cytometers, the imaging throughput is still ~1,000’s cells/s [16] – significantly below the gold-standard non-imaging flow cytometers ~100,000 cells/s.

By contrast, optical time-stretch imaging offers a unique speed advantage over the conventional camera technologies, by achieving continuous frame rate of megahertz or beyond in real-time for high-throughput single-cell imaging in flow [17–21]. Further empowered by interferometry, this technology enables ultrafast QPI for label-free, cancer cell classification [22, 23]. However, revealing sub-cellular features remains challenging in interferometric time-stretch QPI. The major barrier is the exceedingly high group velocity dispersion (GVD) required for achieving diffraction-limited imaging resolution that inevitably comes at the cost of severe optical loss and thus image signal-to-noise ratio (SNR) [24, 25]. Consequently, the degraded SNR lowers the yield of phase-image retrieval and hinders sub-cellular resolution in ultra-large-population single-cell time-stretch QPI (practically >10^5^ cells), which has yet been reported so far [23, 26].

Here we present a new single-cell QPI platform, coined *multiplexed asymmetric-detection time-stretch optical microscopy* (multi-ATOM) that addresses these challenges and thus fills the gap for achieving large-population single-cell biophysical phenotyping at sub-cellular resolution. We demonstrate that multi-ATOM can capture sub-cellular-resolution single-cell QPI images at an ultrahigh-throughput in real-time, at least two orders-of-magnitude faster than classical QPI. Multi-ATOM also outperforms the existing time-stretch QPI modalities in three critical aspects: (i) Bypassing interferometry, multi-ATOM features with ultrafast multiplexed phase-gradient image encoding without excessive GVD and thus loss for achieving sub-cellular QPI resolution; and (ii) multi-ATOM relies on intensity-only phase retrieval algorithm critically guarantees high yield of quantitative phase reconstruction and thus robust single-cell biophysical phenotyping; (iii) Thanks to its sub-cellular resolution preserved at high-throughput, multi-ATOM enables exploration of an additional image contrast, namely dry-mass-density contrast (DC) map, that is sensitive to the *local* variation of dry-mass density (DMD) within the cell and could allow subcellular biophysical single-cell analysis at large-scale. As a proof-of-concept demonstration, we show that multi-ATOM provides sufficient label-free discrimination power for distinguishing two human breast cancer sub-types. It also allows ultralarge-scale label-free image-based classification of human leukemic cells and human peripheral blood monocular cells (PBMCs) (>700,000 cells) at high accuracy (>94%). We therefore anticipate multi-ATOM could find its immediate potential utilities in label-free cancer cell screening in blood.

## 2. Results and Discussions

### 2.1. Working Principles of multi-ATOM

Multi-ATOM leverages the concept of optical time-stretch imaging which implements all-optical image encoding in and retrieval from the broadband laser pulses at an ultrafast line-scan rate (equivalent to the laser repetition rate, i.e. 11.8 MHz in our case) (**Figure 1a**). It is achieved by two image mapping steps: wavelength–time mapping (i.e. time-stretch process) through GVD of light in a dispersive fiber and wavelength–space mapping (i.e. spectral-encoding process) done by spatial dispersion of light using a diffraction grating [17, 20]. Effectively, these two steps enable all-optical laser line-scanning imaging in which the image contrast at each imaged position is encoded with different wavelength within the spectrum of the laser pulse. Here the cells are in a unidirectional microfluidic flow orthogonal to the line-scans which can be digitally stacked to form a two-dimensional (2D) image. In contrast to conventional time-stretch QPI, the distinct feature of multi-ATOM is its ability to encode the local phase gradients of the cells – a quantitative image contrast that have largely been overlooked in the existing time-stretch imaging modalities and yet can be harnessed to obtain the *quantitative phase* of cells without interferometry.

**Figure 1.**
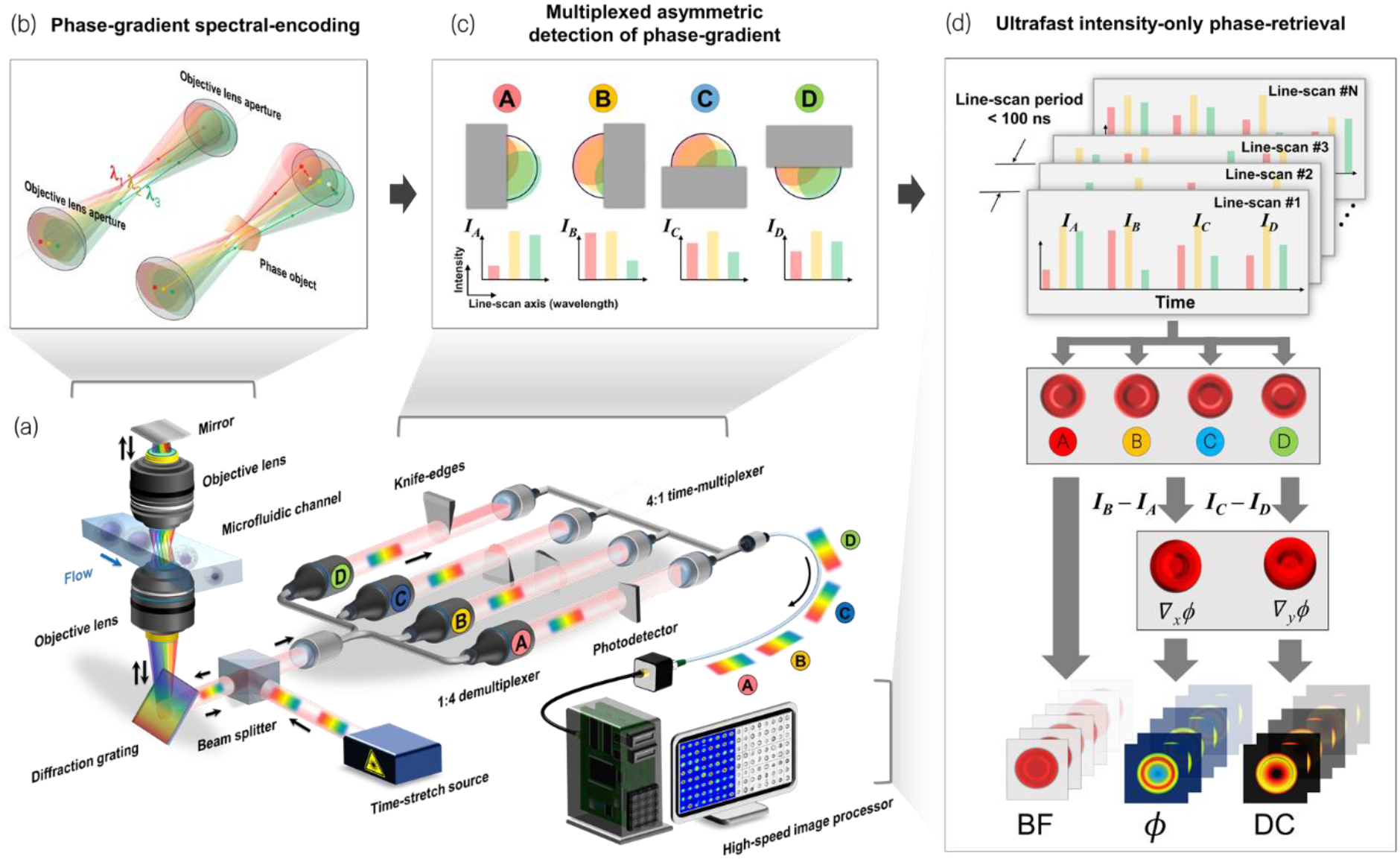
Working principle of multi-ATOM. **(a) System configuration:** The ultrafast line-scan pulsed beam is de-multiplexed into four replicas (A-D) after spectrally encoding the spatial phase-gradient of the flowing cells (flow rate > m/s). Each of these replicas is half-blocked by a knife-edge, followed by time-multiplexing using the fiber delay-lines and realtime signal capture, digitization and processing. **(b) Phase gradient spectral-encoding:** Each minimum spectrally-resolvable beams of the line-scan (i.e. beam *λ*_1_, *λ*_2_, and *λ*_3_, each representing different wavelength) is tilted according to the local phase gradient of phase object (i.e. cell) and is thus partially collected by the objective lens. **(c) Multiplexed asymmetric detection of phase-gradient:** Partial beam-blocking of each beam replica (A-D) from four cardinal directions results in four different transmitted intensity profiles along the spectrally-encoded line-scan, which are used to decode the magnitude and direction of the 2D local phase gradient. **(d) Ultrafast intensity-only phase-retrieval:** the image-encoded line scans (each contains four time-multiplexed replicas) is digitally stacked to form four 2D asymmetrically-detected images (A-D) and thus bright-field (BF) and two phase-gradient images (*∇ϕ_x_* and *∇ϕ_y_*), from which quantitative-phase (*ϕ*) images and dry-mass-density contrast (DC) maps are reconstructed.

Under the condition that spectrally-encoded line-scan illumination is temporally incoherent, the phase gradient and thus quantitative phase across the cell can simply be modelled using geometrical optics (See **Materials and Methods** for details). This model rests on the fact that the local phase gradient of the phase object (i.e. cells) leads to wave-front tilt for each minimum spectrally-resolvable spots of the line-scan and thus partial light collection by the imaging objective lenses (**Figure 1b and S1**). Such light tilt from the image plane is equivalent to a transverse displacement of the minimum spectrally resolvable beam in far-field or on the Fourier plane. In essence, such displacement can be inferred by the fractional intensity loss of each resolvable beam. Hence, the magnitude variation in the phase gradient of the cells along the line-scan, which is spectrally encoded in a single-shot pulse, can be linked to the measured intensity profile across the spectrum.

To evaluate both the magnitude and direction of the 2D local phase gradient, multiple independent measurements of the intensity losses due to light tilt along the x- and y-directions are required. It is easily done by splitting the encoded pulsed-beam into four replicas and partially blocking them by the Foucault-knife-edge method from four cardinal directions before being coupled into four optical fibers (**Figure 1c**). The partial beam block effectively results in an asymmetric beam coupling into the four fibers, which capture the intensities, *I_A_*(*x,y*), *I_B_*(*x,y*), *I_C_*(*x,y*) and *I_D_*(*x,y*) that are linked to the phase-gradient along the four cardinal directions. We previously adopted a similar concept for enhancing the image contrast of label-free time-stretch imaging. [19] Quantifying the phase gradient contrast for time-stretch QPI has not been demonstrated until now. Here, multi-ATOM goes beyond by exploiting multiple sets of spectrally-encoded phase-gradient contrasts to retrieve the quantitative phase. More precisely, the phase-gradients along the *x*- and *y*-directions (∇*ϕ_x_* and ∇*ϕ_y_*) are linked to the normalized intensity losses *R_x_*(*x,y*) and *R_y_*(*x,y*), which are proportional to the difference images, i.e. *I_B_* − *I_A_* and *I_C_* – *I_D_* respectively (**Figure 1d and 2a**, and see **Materials and Methods**). The quantitative phase image, *ϕ*, (**Figure 1d and 2a**) from the phase gradients is evaluated by complex Fourier integration with a calibration factor obtained through imaging a known quantitative phase target. Note that only the intensities of the four images are needed and no iterative process is involved for phase retrieval. Other than quantitative phase, the bright-field contrast, i.e. BF (**Figure 1d**) can also be obtained from the four asymmetrically-detected signals (See **Materials and Methods**). To ensure ultrafast operation, four replicas are captured by a time-multiplexing scheme: replicas are firstly coupled into four individual fiber delay lines, each with different length, and then recombined by a multiplexer to become temporally separated. Eventually, these replicas are captured within each single-shot line-scan period (~84 ns) to achieve ultrafast QPI (**Figure 1d and 2a**).

**Figure 2.**
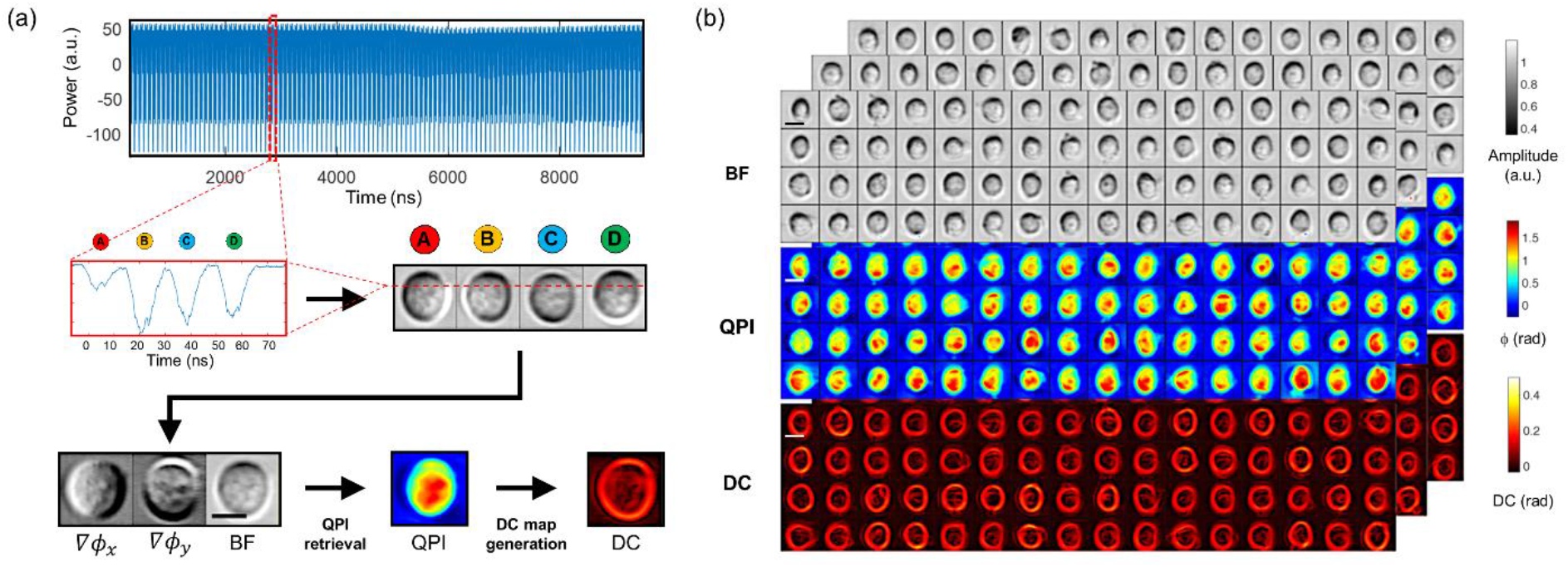
Image reconstruction pipeline illustration and representative cell images. **(a) Image reconstruction:** The continuous data stream is first truncated into data segments that each records a single cell. A single segment consists of a few tens of line scan, within each four pulse replicas are time-multiplexed (zoom-in windows). By reshaping the segment, an image including four asymmetric-detection contrast sub-images from the same single cell can be obtained (A-D) from which two phase-gradient-constrast (*∇ϕ_x_* and *∇ϕ_y_*) and a bright-field contrast (BF) images are arithmatically combined. Based upon, a quantitative phase image (*ϕ*) is then retrieved by complex Fourier integration as well as the dry-mass-density contrast (DC) image. **(b) Representative multi-contrast multi-ATOM cell images:** BF, QPI and DC single-cell images of leukeimic cells (THP-1) which are captured at a throughput of >10,000 cell/sec. Scale bar = 10μm.

We can derive a multitude of image features, as the biophysical phenotypes of the cells, from both the quantitative phase and BF images. For instance, we extract the typical bulk parameters, e.g. cell volume, roundness, optical opacity from the BF image whereas averaged dry mass (DM) and DMD from the phase image (See **Table S1** for detailed derivation). Beyond that, taking the advantage of the sub-cellular resolution preserved at high-throughput in multi-ATOM, we further transform each quantitative phase image into a *DC map* which visualizes and quantifies the *local* variation of DMD across the cell (**Figure 2a and S2** and **Table S1**). We can thus extract additional sub-cellular resolvable phenotypes by quantifying the statistics of local DC of individual cells, such as DC1, DC2 and DC3 as the 1^st^, 2^nd^ and 3^rd^ order statistical moments of DC which relate to the variation and the skewness of *local* DC distribution respectively (**Table S1**). We note that the statistics of DC values across the cell is thus different from the *global* phase statistics obtained directly from the phase image [5]. Altogether, multi-ATOM enables label-free single-cell biophysical phenotypic profiling within an enormous population of cells, as discussed in later section.

### 2.2 Imaging performance of multi-ATOM

We evaluated the basic performance of multi-ATOM in terms of its resolution, phase accuracy and stability. From the retrieved phase image of the phase target, the height profile of the USAF resolution chart target was measured to be ~100 nm, consistent to the target specification (**Figure S3a-b**). The corresponding *QPI* spatial resolution was also measured as 1.38 μm and 0.98 μm along the spectral encoding (fast-axis) and sample scanning (slow-axis) direction respectively, according to the smallest features resolved by multi-ATOM (i.e. group 8 element 4 and group 9 element 1 respectively in **Figure S3a-b**). On the other hand, the shot-to-shot fluctuations and the long-term stability of the measured optical path lengths (derived from quantitative phase) over 4 hours were measured to be 4.5 nm (8.8 mrad) and 8.3 nm (16.2 mrad) respectively (**Figure S3d**). It demonstrates the high stability of multi-ATOM, contributed by its interferometry-free measurement which is less sensitive to ambient mechanical perturbation. We next characterized the cell imaging performance of multi-ATOM in a microfluidic flow at an imaging throughput of >10,000 cells/sec (**Figure 2a-b**). Multi-ATOM clearly shows the ability to capture motion-blur-free amplitude, phase-gradient, phase, and DC images of individual fast-flowing cells (2.3 m/s) and reveal their sub-cellular textures at an ultrafast imaging rate (a line-scan rate of 11.8 MHz) – a combined capability absent in both conventional camera-based QPI and interferometric time-stretch QPI (**Figure 2b**).

Achieving high spatial resolution in any time-stretch imaging modalities is often complicated by the interplay among GVD, optical loss, and data sampling rate [25]. Notably, exceedingly high sampling rate is required to reduce GVD and dispersive loss. But this condition can only be achieved in the high-end oscilloscopes (typically >40 GSa/s) at the expense of high-throughput continuous single-cell image capture in real-time. On the other hand, using the system with lower sampling rate (e.g. field programmable gate array (FPGA) platform typically at <10 GSa/s) could improve the throughput performance at the cost of considerably higher GVD required for achieving diffraction-limited resolution [27]. The GVD requirement becomes even more stringent in interferometric time-stretch QPI (or interferometric time-stretch microscopy (iTM) [24]) where extra GVD is required to ensure high fringe visibility of the spectrally-encoded interferogram, and thus diffraction-limited QPI. However, such an excessive GVD inevitably results in significant optical loss, and thus degrades the imaging sensitivity.

In contrast, multi-ATOM bypasses the use of interferometry. This offers the key advantages of mitigating the stringent demand for high GVD and preserving sub-cellular resolution at high imaging throughput. To demonstrate such significance, we configured an integrated system, which enables *simultaneous* multi-ATOM and iTM of the *identical* cells under a diffraction-limited resolution condition (**Figure S4 and Section S1.2 in Appendix S1**), and evaluated the effect of GVD on the performance in the two modalities (**Figure 3** and **Materials and Methods**). According to time-stretch theory, changing GVD at a fixed sampling rate has the equivalent effect on image resolution to varying the sampling rate at a fixed GVD [27]. Hence, here we study the GVD effect at the fixed sampling rate adopted in this work (4 GSa/s) by defining a range of equivalent GVD (GVD_*eq*_), which is governed by the inverse relationship between GVD and sampling rate (See the detailed theory in **Materials and Methods**).

**Figure 3.**
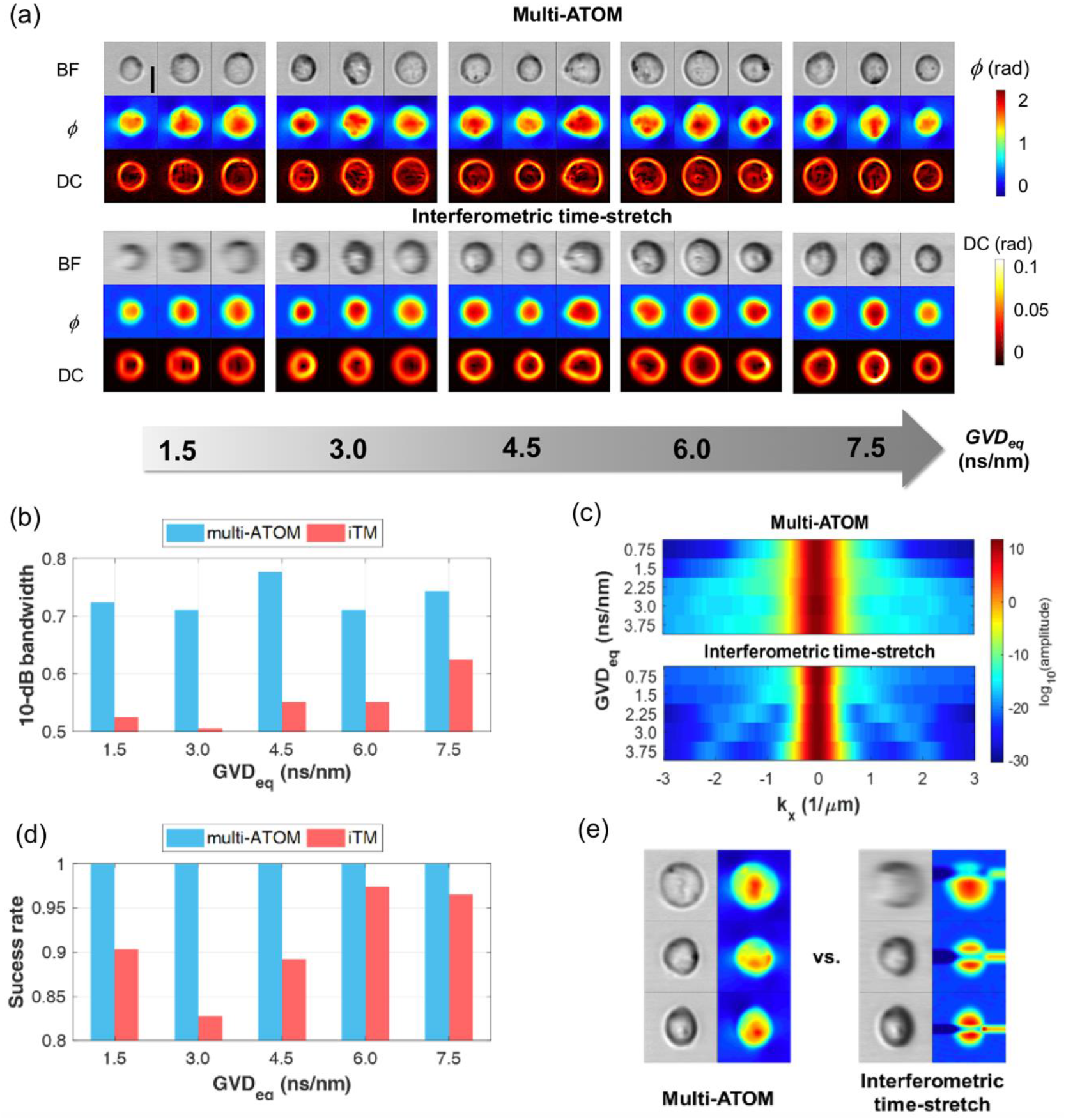
Performance comparison between multi-ATOM and interferometric time-stretch microscopy (iTM). **(a) Image quality:** Each cell is simultaneously imaged by multi-ATOM and iTM methods. The reconstructed images of each cells, with bright-field (BF), quantitative phase (ϕ) and dry-mass-density contrast (DC) image contrasts, are compared under 5 different equivalent group velocity dispersion (GVD_eq_) conditions (**Materials and Methods**), **(b-c) Spectral bandwidth analysis:** (b) Averaged 10-dB bandwidth of ϕ images (along the fast axis) and (c) the corresponding spectra obtained from both systems at different GVDeq. (c). Colorbar: logarithmic amplitude. **(d-e) Phase retrieval yield analysis:** (d) Comparison of success rate of phase retrieval between multi-ATOM and iTM based on captured images of ~4600 cells and (e) three representative failed cases of phase retrieval of iTM.

In general, except the overall cell shape is visualized, the sub-cellular texture is lost in iTM images as compared with that of multi-ATOM (**Figure 3a**). Although the resolution of iTM increases with *GVD_eq_*, which can be quantified in the corresponding signal bandwidth, the resolution in multi-ATOM generally remains higher than that in iTM (**Figure 3b-c**). According to time-stretch imaging resolution analysis [21, 25, 28], at least two times higher in GVD is required in iTM (2.29 ns/nm) compared to multi-ATOM (1.14 ns/nm), in order to achieve similar imaging resolution (**Figure S5 in Appendix S1**). The resolution advantage of multi-ATOM compared to iTM becomes more obvious when we further transform the quantitative phase image into a 2D DC map, which reveals more clearly the sub-cellular structures using multi-ATOM (**Figure 3a**).

Another benefit from the interferometry-free operation is that the quantitative phase retrieval in multi-ATOM bypasses the need for the iterative phase unwrapping algorithms commonly used in QPI [29–31]. Not only can it reduce the computational complexity which favours ultrafast real-time image processing, but also more importantly, eliminates the phase unwrapping error which is susceptible to noise and low fringe visibility [22, 29]. Hence, phase retrieval in multi-ATOM is essentially error-free compared to that in iTM (**Figure 3d-e**). Note that the reduction in success rate in iTM (i.e. ~5-15%), may not be tolerable in view of practical detection of small populations in an ultra-large-scale analysis of heterogeneous cell population, not to mention the even worse success rate in reality because of an extra 10-dB degradation of SNR accompanied with doubling the GVD for achieving sub-cellular resolution.

### 2.3 Single-cell image-based biophysical phenotyping and classification using multi-ATOM

We investigated the ability of multi-ATOM-derived single-cell biophysical phenotypes, especially the sub-cellularly resolvable features, to discriminate two breast cancer lines, MCF-7 and MDA-MB-231 [32], which represents two sub-types of breast carcinoma (**Figure 4a**). Both the DC parameter (i.e. DC1) and opacity are found effective, as label-free biophysical markers, to distinguish the two cell types in a mixed population (**Figure 4b** and see also the verification of the pure populations shown in **Figure S6**). In contrast, such mixture of two populations are not resolvable in standard flow cytometry based on scatter light measurements (inset of **Figure 4b**) – signifying the significance of quantitative label-free, high-resolution imaging for cytometry applications.

**Figure 4.**
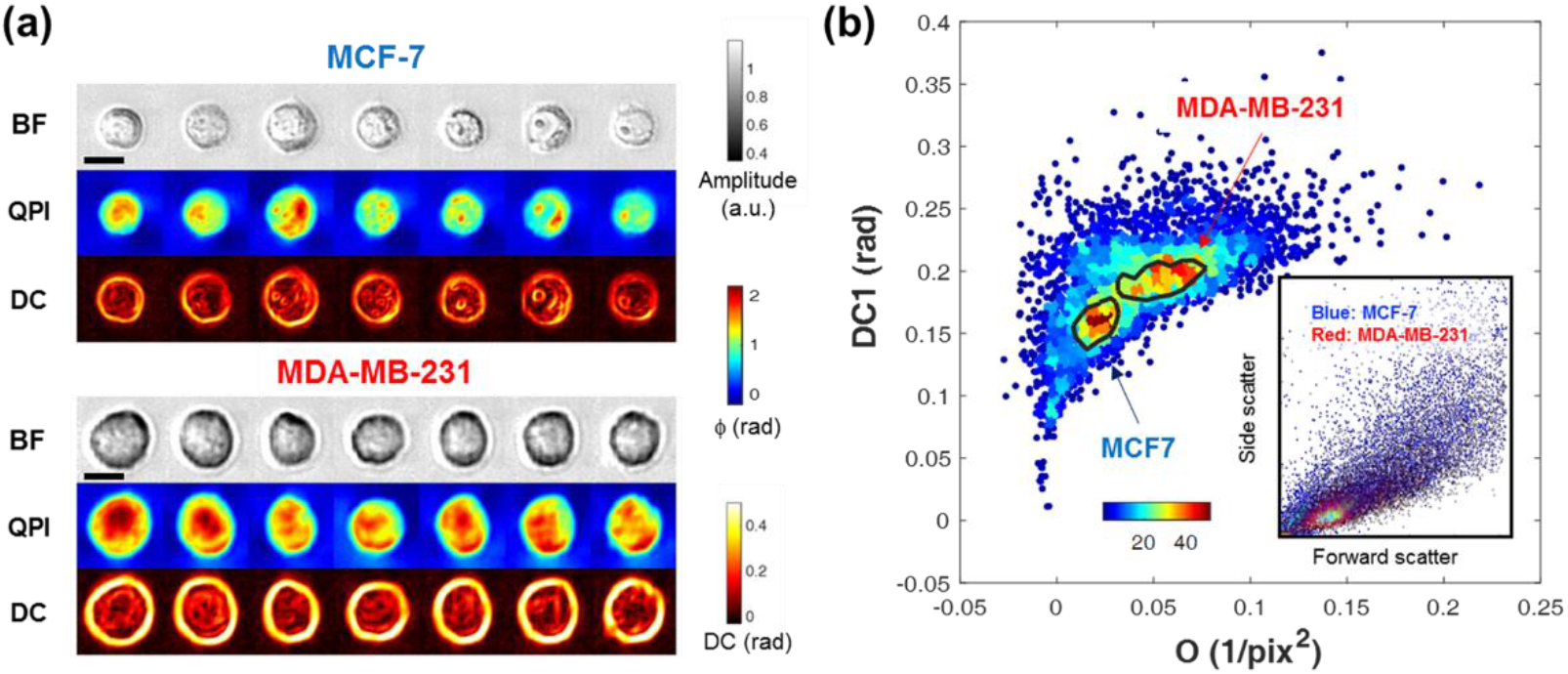
Label-free classification of two breast cancer cell types. **(a) Representative cell images:** BF, QPI and DC single-cell images of two types of breast cancer cells (top: MCF-7, bottom: MDA-MB-231). Scale bar = 10μm. **(b) Discrimation power of multi-ATOM-derived features:** A bivarate density plot from multi-ATOM showing distinctive distributions for two cancer cell types with the 60%-density-contours overlaid, while that of converntional flow cytometer (inset) shows an highly overlaped distribution. Color bar: local density of data points.

We further assessed the high-throughput capability of multi-ATOM in ultra-large-scale singlecell imaging (more than 700,000 cells). The underlying rationale is that the combined advantage of high-throughput and high-resolution imaging brought by multi-ATOM could provide a sufficient statistical power in single-cell biophysical phenotyping, which was once regarded to lack the specificity for cell-based assay, to reveal the diversity of cell populations. This is evident from the heterogeneity revealed in the bivariate plots (sub-cellular texture phenotype DC2 versus volume) of both cell types, especially the PBMC sample, which is known to be highly heterogeneous (**Figure 5a-b**). Precisely because of such diversity, high-dimensional biophysical phenotypic analysis is necessary to ensure cell classification with sufficiently high sensitivity and specificity. It is challenging to visually distinguish the heterogeneous clusters of two cell types until at least three biophysical markers are used (**Figure 5c**). Our receiver-operating-characteristic (ROC) analysis shows that using 12 multi-ATOM-derived biophysical phenotypes (**Table S1**), notably the QPI-related markers (e.g. DMD, DC2, and DC3), further improves the classification sensitivity and specificity (THP-1 versus PBMC), resulting in an area-under-curve (AUC) as high as 0.972 (**Figure 5d**). These results demonstrate that multi-ATOM enables high-resolution QPI at ultrahigh throughput – a capability absent in both conventional camera-based QPI and interferometric time-stretch QPI. It thus allows robust analysis of the high-dimensional, label-free, spatially-resolved biophysical single-cell data at large-scale.

**Figure 5.**
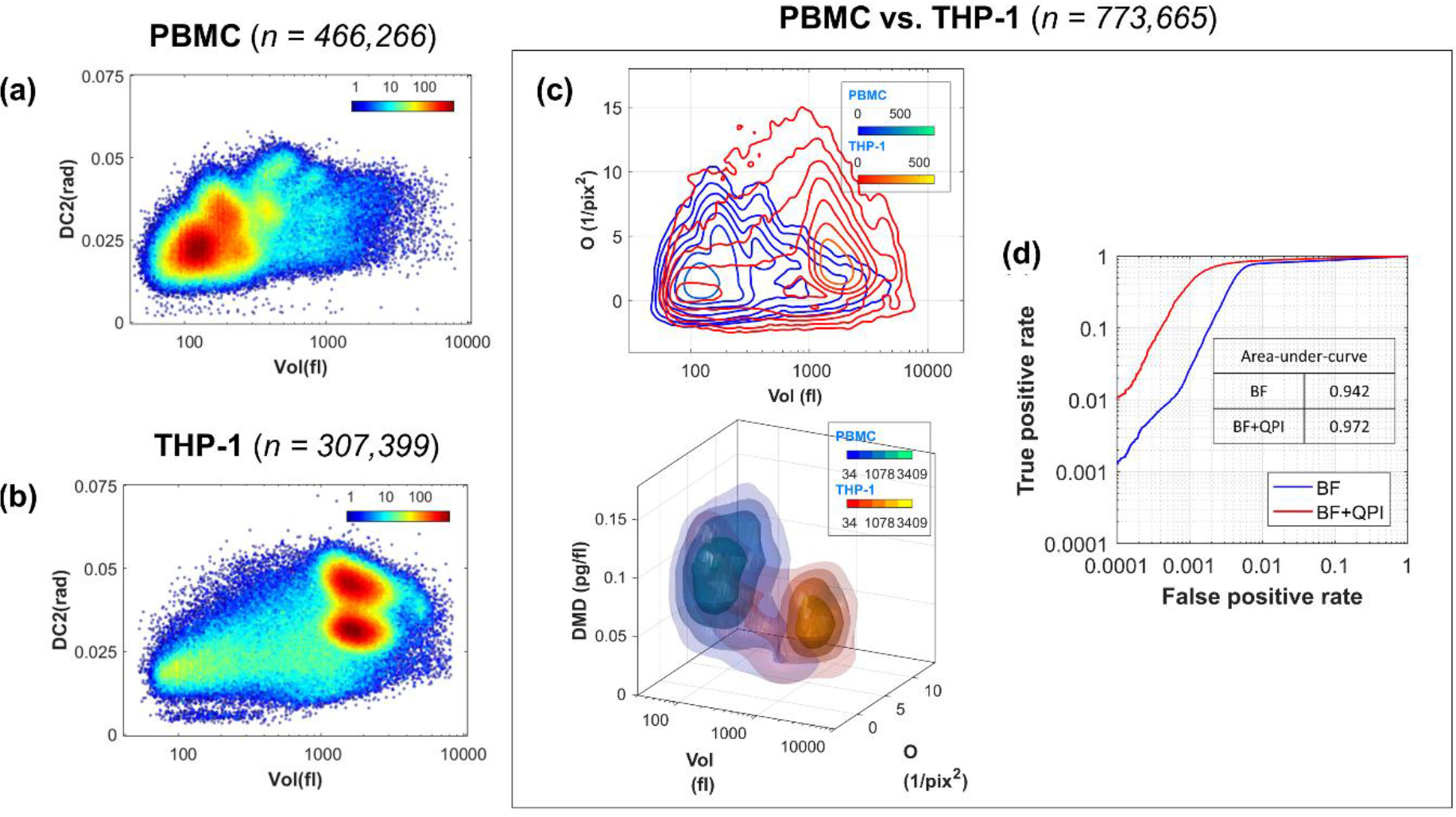
Ultra-large-scale label-free multivariate classification of cancer cells and blood cells. Both **(a)** peripheral blood mononuclear cells (PBMC, n = 466,266) and **(b)** leukemic monocytes (THP-1, n = 307,399) show high degree of heterogeneity, as clearly revealed in the bivariate color-coded density plots. **(c)** (top) A contour (2D) map and (bottom) an iso-surface (3D) map of the two cell types. **(d)** The receiver-operating-characteristic (ROC) curves showing the label-free classification accuracy. The significance of including QPI-image-derived features on classification accuracy is reflected by the increased area-under-curve (AUC).

## 3. Conclusion

Based upon an ultrafast, interferometry-free, phase-gradient encoding concept, multi-ATOM has an overall strength of enabling ultrahigh throughput QPI without compromising the sub-cellular resolution – a capability that is absent in both conventional camera-based QPI and interferometric time-stretch QPI. We demonstrated that multi-ATOM, at least two orders-of-magnitude faster than the classical QPI approaches, can efficiently reconstruct individual sub-cellularly-resolvable multi-contrast, label-free single-cell images (both the phase, amplitude and DC images) within an enormous population (>700,000 cells). We showed that *both* ultrahigh imaging throughput and high label-free sub-cellular content realized in multi-ATOM critically enable high statistical discrimination power to classify leukemic cells and PBMCs. Such capability is generally challenging with current biophysical phenotyping techniques [4, 23, 33, 34]. Therefore, this label-free single-cell imaging technique could find new ground in many applications, notably new cost-effective liquid biopsy strategies, e.g. and circulation tumor cell detection and pre-screening, in which number of known biochemical markers and thus the bias and the cost of assay can be minimized; and new form of SCA in which the catalogue of biophysical markers could complement and correlate the cellular heterogeneity information provided by the assays based on biochemical markers, and thus enable strategic cost-effective ultra-large-scale SCA.

## 4. Materials and Methods

### 4.1. Multi-ATOM system configuration

The system consists of three functional modules (**Figure 1a**): (1) a time-stretch module for wavelength-time mapping, (2) a spectrally-encoded imaging module for wavelength-space mapping and (3) an asymmetric-detection module for phase-gradient encoding/decoding. In the time-stretch module, a home-built all-normal dispersion (ANDi) laser (centre wavelength = 1064 nm, bandwidth = 10 nm and repetition rate = 11.8 MHz) generates the pulse train (pulse width = ~12 ps), which is then stretched in time by a 10-km single-mode fiber (a total GVD = 1.78 ns/nm and 17 dB loss at 1064 nm (OFS, US)) and optically amplified by a ytterbium-doped fiber amplifier module (output power = 36 dBm, on-off power gain = 36 dB) – resulting in an all-optical wavelength-swept source for multi-ATOM. (2) In the spectrally-encoded imaging module, a diffraction grating (groove density = 1200/mm, Littrow angle = 42.3) spatially disperses the time-stretched pulses to generate a one-dimensional (1D) spectral shower. A pair of objective lenses (numerical aperture (NA) = 0.75 (front) and 0.8 (back)) forms a double-pass configuration for imaging the suspended cells flowing in a custom-designed microfluidic chip (flow rate > 1 m/s) along the orthogonal direction to the spectral shower. The same grating recombines the reflected image-encoded spectral shower into a single collimated beam. Note that the local phase-gradient induced by the imaged cells is spectrally encoded into intensity-loss of each minimally resolvable point across the spectral shower. (3) In the asymmetric-detection module, a one-to-four de-multiplexer splits the image-encoded pulsed beam into four identical replicas, each of which is half-blocked by a knife-edge from four primary cardinal directions (i.e. right, left, top and bottom) (**Figure 1a,c**). Then, the four beams are time-multiplexed by the fiber delay-lines and a fiber-based four-to-one multiplexer. Hence, the four image-encoded pulses (all correspond to the same line scan) are separated in time, i.e. time-multiplexed. A high-speed single-pixel photodetector (electrical bandwidth = 12 GHz (Newport, US)) is used for continuous signal detection. For digitization, a real-time field programmable gate array (FPGA) based signal processing system is used (electrical bandwidth = 2 GHz, sampling rate = 4 GSa/s, throughput >10,000 cell/s). Custom logic was implemented on the FPGA to automatically detect and segment cells from the digitized data stream in real time, at a processing throughput equivalent to >10,000 cell/s. This segmentation step is critical in maximizing the efficiency of the downstream data storage subsystem. All segmented cell images are sent through four 10G Ethernet links and are stored by four data-receiving hosts with a total memory capacity of over 800 GB that can be extended easily.

### 4.2. Phase retrieval algorithm of multi-ATOM

A key prerequisite of robust phase retrieval in multi-ATOM is to create a temporally incoherent spectrally-encoded line-scanning illumination. Under this condition, which can be accomplished with high GVD, the propagation characteristics of each minimum spectrally-resolvable beam across the line-scan can adequately be modelled using geometrical ray-tracing. This condition is satisfied when the time delay between the adjacent minimum spectrally resolvable spots τ_min_ is significantly greater than the effective coherence time τ_c_ (defined by the spectra resolution of the spectrally-encoded illumination, *δλ*), i.e. τ_min_ ≫ τ_c_, where τ_min_ = *δλ* · *GVD* and *τ_c_* = *λ*^2^/*cδλ. c* is the speed of light and *λ* is the centre wavelength of the light. Given that *δλ* = 0.2 nm, and *GVD* =1.78 ns/nm employed in our system, the aforementioned condition is satisfied.

The central idea of phase retrieval of multi-ATOM is integrating the phase gradients encoded in asymmetrically detected signal to obtain the quantitative phase image (*ϕ*(*x,y*)). In detail, two orthogonal phase gradient images (*∇_x_ϕ*(*x,y*) and *∇_y_ϕ*(*x,y*)) are reconstructed from 4 asymmetrically-detected intensity images (*I_A_*(*x,y*), *I_B_*(*x,y*), *I_C_*(*x,y*) and *I_D_*(*x,y*) – A, B, C, D in **Figure 1**) through three mapping equations: mapping from phase gradient to tilt angle of wavefront (Eqn. (1)); mapping from tilt angle of wavefront to transverse beam translation (Eqn. (2)); mapping from transverse beam translation to intensities of the four asymmetrically-detected images (Eqn. (3)). Throughout the derivation, the spatial coordinate of the image is indexed by symbols, *x* and *y*, and a flat beam profile (i.e. Köhler illumination on the frontal plane in **Figure S1**) is considered for simplicity which can be approximated by overfilling the objective aperture.

A phase gradient presents when there is a spatial phase difference due to the refractive index mismatch and/or the thickness profile across phase objects, e.g. biological cells. Consider a focused light beam illuminates on a phase object, refraction of light (i.e. wavefront tilt) occurs because of the local phase gradient, as illustrated in **Figure S1**. In multi-ATOM, this phenomenon applies to each minimally resolvable spectrally-encoded beam along the spectral shower. Assuming the phase gradient is small and smooth across the phase object, one could apply the concept of ray optics to formulate the phase gradient, which is proportional to the tilt angle of wavefront under the paraxial approximation:

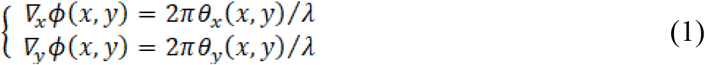

where *∇_x_ϕ* and *∇_y_ϕ* are the phase gradients along the *x* and *y* direction; *θ_x_* and *θ_y_* are the tilt angles of wavefront along x and y direction; *λ* is the center wavelength of light. The light tilt effectively leads to a transverse translation of the beam profile in the frontal plane. From the geometry of the light cone (transverse plane and sagittal plane in **Figure S1**), the translations (*δL_x_* and *δL_y_*) along x and y direction can be related to the corresponding tilt angle of wavefront (*θ_x_* and *θ_y_*) by:

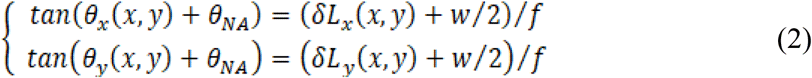

where *θ_NA_* and *f* are the acceptance angle and focal length of the objective lens, respectively. Here we assume the aperture in the frontal plane of the object to be a square with the sides of *w*. For simplicity, the two objective lenses are considered to have the same numerical aperture (NA). The beam translation causes an incomplete light coupling to the back objective such that the power of coupled light decreases (frontal plane in **Figure S1**). Practically, objective lenses are not necessary to have identical NA. Because of the double-pass configuration, the effective NA is always limited by the smaller objective NA – the identical NA situation can be relaxed to be NA_back obj_ ≥ NA_front obj_. In other words, the local light tilt and thus the phase-gradient are spectrally encoded into the power of each minimally resolvable beam across the spectral shower.

To decode the relative power loss due to the beam translations and thus the phase gradients along both the *x* and *y* directions, multiple intensity measurements of the same image are required. We adopt the asymmetric detection scheme that involves partial-beam blocks on four beam replicas from the same line-scan (**Figure 1a**). At each pixel coordinate (x, y), the relationship of the normalized intensity loss along the x and y directions (*R_δx_*(*x, y*) and *R_δy_*(*x,y*)), the intensities of these four image replicas (*I_A_*(*x,y*), *I_B_*(*x, y*), *I_C_*(*x,y*) and *I_D_*(*x,y*)) and the local beam translations (*δL_x_*(*x,y*) and *δL_y_*(*x,y*))) can be expressed by:

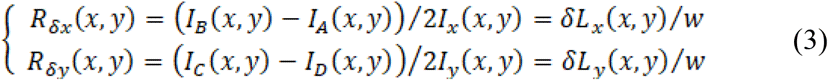

where *I_x_*(*x,y*) and *I_y_*(*x,y*) are absorption-contrast images along x and y direction (**Materials and Methods Section 4.3**).

By combining Eqns. (1) – (3), the two orthogonal phase gradients (*∇_x_ϕ*(*x,y*) and *∇_y_ϕ*(*x,y*)) and thus the complex phase gradient (*∇ϕ*(*x,y*)) can be related to the measured intensity loss ratio along the x and y directions (*R_δx_*(*x,y*) and *R_δy_*(*x, y*)).

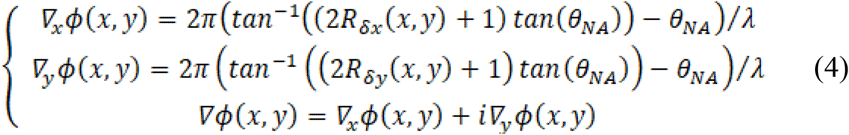

Eqn. (4) indicates that in order to quantify the phase gradient and thus the phase, the only experimental quantity are the two normalized intensity losses (*R_δx_*(*x,y*) and *R_δy_*(*x,y*)). Specifically, they are the intensity ratios of the asymmetrically-detected images along the *x* direction, (i.e. *I_B_*(*x,y*) − *I_A_*(*x,y*)) and the *y* direction (i.e. *I_C_*(*x,y*) − *I_D_*(*x,y*)) to its absorption-contrast images (*I_x_* and *I_y_*) respectively. (**Materials and Methods Section 4.3**).

Complex Fourier integration is then applied on the phase gradient (*∇ϕ*(*x,y*)) to obtain the quantitative phase image (*ϕ*(*x,y*)),

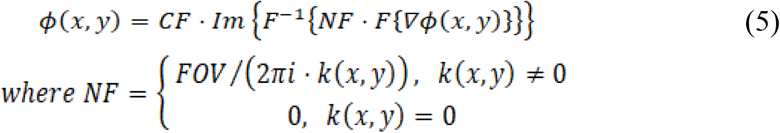

where *Im* is the imaginary part of a complex number; *F* and *F*^−1^ are the forward and inverse Fourier transform operators respectively; *NF* is a normalization factor for quantifying the phase and avoiding singularity in the integration operation; *k*(*x,y*) is the 2D wavenumbers; *FOV* is the 2D field-of-view; *CF* is the calibration factor for correcting the systematic phase deviation arise from non-ideal system setting, i.e. Gaussian beam profile and circular aperture.

### 4.3 Bright-field image reconstruction in multi-ATOM

The four normalized asymmetrically-detected images (*I_A_*(*x,y*), *I_B_*(*x,y*), *I_C_*(*x,y*) and *I_D_*(*x,y*) - A, B, C, D in **Figure 1c**) are divided into two sets of images based on their asymmetric detection orientations, i.e. *I_A_*(*x,y*) & *I_B_*(*x,y*) as the horizontally blocked images, and *I_C_*(*x,y*) & *I_D_*(*x,y*) as the vertically blocked images. In each set, the intensity values of two images are compared pixel-by-pixel (i.e. *I_A_*(*x,y*) vs. *I_B_*(*x,y*) and *I_C_*(*x,y*) vs. *I_D_*(*x,y*)) such that the pixels with maximum intensity from one of the two images are selected to form the absorption-contrast image of each set (*I_x_*(*x,y*) and *I_y_*(*x, y*)).

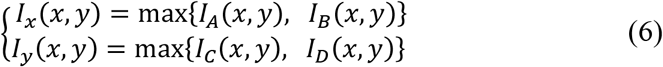

The bright-field image of a cell (*BF*(*x,y*)) is the sum of two resulting absorption contrasted images (*I_x_*(*x,y*) and *I_y_*(*x,y*)).

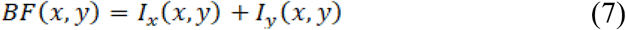

From the above image reconstruction algorithm, the scattering loss caused by the cells would be minimized, such that the bright-field image would represent mostly the absorption loss.

### 4.4 Reconstruction of dry-mass-density contrast (DC) images

A DC image is transformed from a quantitative phase image through 2D convolution with a diffraction-limited-spot-size kernel within which statistical standard deviation of the phase value is calculated (**Table S1**). As a result, quantitative phases are transformed to DC values at each pixel. A binary mask obtained from the corresponding bright-field image is applied to the DC image to isolate the cell body from the whole image. A statistical histogram of DC of the cell is then generated from the masked DC image. The standard deviation and the skewness, which corresponds to DC2 and DC3 respectively, are calculated from the histogram (**Table S1**).

### 4.5. Human peripheral blood mononuclear cell (PBMC) isolation from human buffy coat

Human buffy coat was provided by the Hong Kong Red Cross; and all the blood donors have given written consent for clinical care and research purposes. The research protocol is approved by Institutional Review Board at the University of Hong Kong. 2 mL of human buffy coat/fresh blood mixed well with 2mL of phosphate-buffed saline (PBS) and then carefully layered on top of 2 mL of Ficoll (GE Healthcare Life Sciences, US) inside a 10ml centrifuge tube without mixing. The centrifuge tube containing the above-mentioned solutions was transferred to centrifugation under 400xg for 20 minutes after which the fluid is separated into five layers – corresponding to plasma, PBMCs, Ficoll, granulocytes and RBCs (from top to bottom). PBMCs were carefully pipetted out from the corresponding layer and transferred to separated centrifuge tubes. The extracted PBMCs were then centrifuged and re-suspended with 1x PBS to achieve cell count of ~10^5^ cells per mL.

### 4.6. Sample preparation of cell lines

Culture medium for THP-1 (TIB-202™, ATCC, US), culturing was formulated with 90% Roswell Park Memorial Institute (RPMI) -1640 medium (Thermo Fisher Scientific, US), 10% fetal bovine serum (FBS) and 1% 100x antibiotic-antimycotic (Anti-Anti, Thermo Fisher Scientific, US). Culture medium for MDA-MB-231 (HTB-26^™^, ATCC, US), and MCF-7 (HTB-22D™, ATCC, US) culturing in DMEM medium (Gibco^−^) supplemented with 10% PBS and 1% 100x antibiotic-antimycotic (Anti-Anti, Thermo Fisher Scientific, US). Cells were cultured in a 5% CO2 incubator under 37 °C and the medium was renewed twice a week. Cells were pipetted out adjusted to be around 10^5^ cells per mL of 1x PBS. Prevention of mycoplasma contamination was done by adding Antibiotic-Antimycotic (Thermo Fisher Scientific, US) during cell culture. Cellular morphology was routinely checked during cell culture under light microscope prior to imaging experiments

## 6. Data sharing and data accessibility

Data is available upon request.

## 7. Acknowledgements funding

The work is partially supported by the Research Grants Council of the Hong Kong Special Administrative Region of China (HKU 17207715, 17207714, HKU 720112E, HKU 719813E, T12-708/12-N, C7047-16G), Innovation and Technology Support Programme (ITS/090/14, GHP/024/16GD), and the University Development Funds of the University of Hong Kong.

## Graphical Abstract

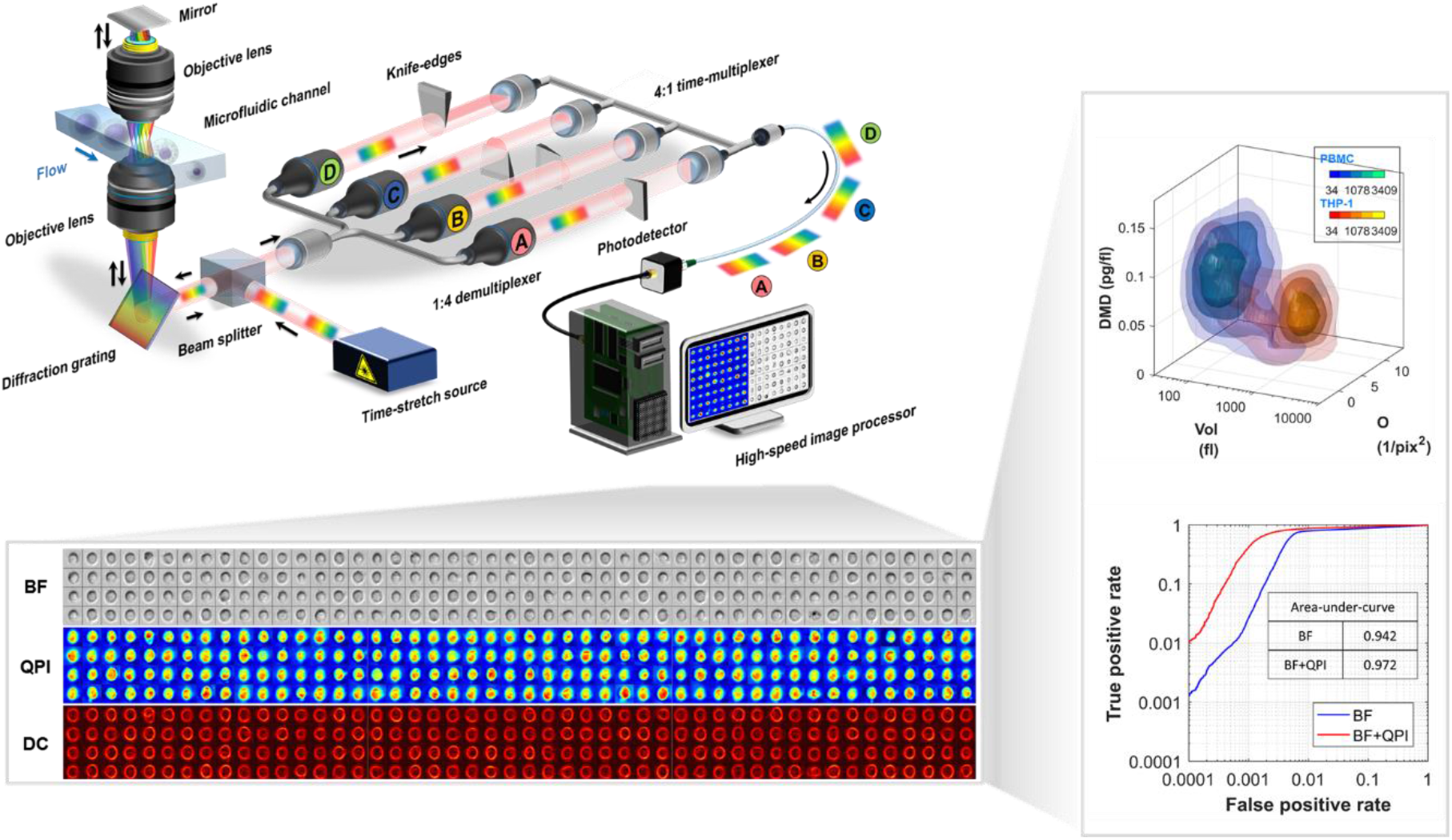

An ultrafast quantitative phase imaging (QPI) modality, called multi-ATOM, is reported that enables ultralarge-scale, multi-contrast, label-free single-cell imaging without compromising subcellular resolution. This capability, otherwise challenging in current QPI, critically enable high label-free statistical discrimination power to allow multi-variate cell-type classification at high accuracy (> 94%), suggesting its potential to empower new strategies in large-scale biophysical single-cell analysis with applications in biology and enriching disease diagnostics.

