## Supplementary files for "Multi-ATOM: Ultrahigh-throughput single-cell quantitative phase imaging with subcellular resolution"

**S1. Supplementary Methods**

**S1.1. Image resolution evaluation of multi-ATOM and iTM**

The overall image resolution of multi-ATOM and iTM along the spectral shower (x-axis) and along the flowing direction (y-axis) are governed by different factors:

Resolution along spectral shower (x-axis)

- Multi-ATOM: Multi-ATOM performs imaging through a 2-step information mapping across the space, wavelength, and time domains, namely wavelength-space and wavelength-time mapping. As a result, the imaging resolution of multi-ATOM is not simply governed by diffraction, and is instead influenced by 3 factors relevant to the time-stretch process: (i) spatial dispersion, which relates to the spectral resolution of the diffraction grating and the diffraction limit, (ii) stationary-phase-approximation (SPA), which describes the ambiguity of the wavelength-time mapping and (iii) the bandwidth of digitizer, which defines the temporal resolution of the real-time signal acquisition [1]. The overall resolution of multi-ATOM is thus limited by the minimal resolution among these 3 factors, i.e. the largest value of the smallest resolvable points. The smallest resolvable point in each of these regimes is respectively evaluated as: (i) 1.31 μm, (ii) 0.39 μm and (iii) 0.84 μm, showing that, the image resolution of multi-ATOM along spectral shower is limited by diffraction, i.e. 1.31 μm.
- iTM: With an additional interferogram, iTM requires at least a double of sample points to resolve the same resolution as muli-ATOM under the same GVD. Therefore, the smallest resolvable points in three domains are evaluated as: (i) 1.31 μm, (ii) 0.78 μm and (iii) 1.69 μm. It shows that iTM can only achieve detector-limited resolution, 1.69 μm.

Resolution along flow direction (y-axis)

- Multi-ATOM: As there is no spectral encoding involved in the flow direction, the image resolution of multi-ATOM is diffraction limited, i.e. 0.87 μm, following the Rayleigh criterion.
- iTM: Same as multi-ATOM (0.87 μm) as there is no spectral encoding in the flow direction.

**S1.2 Experimental configuration for performance comparison between iTM and multi-ATOM**

To compare the imaging performance between iTM and multi-ATOM in a continuous data capturing scenario, we introduced two key modifications in the multi-ATOM system (**Figure S4**). Firstly, to capture both iTM and multi-ATOM image data of the same cell simultaneously, an additional interferometry module was added. Specifically, a 50:50 beam splitter (Thorlabs, US) was placed right before the asymmetric-detection module to equally split the beam into two replicas. One of the replicas entered the multi-ATOM module; another enters the iTM module and was interfered with a reference beam (with the same power) using another 50:50 beam splitter which helped maximizing the interference fringe visibility. The resultant interferogram was detected by another single-pixel photodetector (electrical bandwidth = 8.5 GHz, (Newport, US)). Secondly, a real-time oscilloscope with 4-times larger in detection bandwidth than the FPGA module (electrical bandwidth = 16.8 GHz, sampling rate = 80 GSa/s (Keysight, US)) was employed. Note that the real-time high-bandwidth oscilloscope was used here to ensure the raw data from both iTM (temporal interferogram) and multi-ATOM (time-multiplexed replica) were sufficiently oversampled such that diffraction-limited resolution was achieved in both multi-ATOM and iTM at the GVD given by the dispersive fiber available in the present work, i.e. GVD = 1.78 ns/nm. The fringe density was configured (by tuning the reference arm length in the interferometer) to be sufficiently high to achieve the diffraction-limited resolution and able to be resolved at the given GVD and sampling rate. These are the necessary conditions for fair comparison of image quality between multi-ATOM and iTM (see the next section). We note that in practice, the oscilloscope is incompatible with continuous cell image capture at high-throughput because of its limited memory depth.

**S1.3 Data processing pipeline (multi-ATOM & iTM)**

The data processing pipeline in this case (i.e. **Figure 2**) can be divided into 3 main tasks: (1) mask generation, (2) multi-ATOM data processing and (3) iTM data processing.

To ensure a fair comparison between multi-ATOM and iTM, especially in terms of feature extraction, a binary mask is first computed from multi-ATOM data for each of the cell which will be applied to both multi-ATOM and iTM images of the same cell in the next two tasks. The systematic spatial shifts between the mask and the cell bodies in multi-ATOM/iTM are also estimated here such that there will be no time difference in searching for cell body while applying the mask. The mask and the corresponding shifts are then recorded.

After the mask generation, the multi-ATOM data is processed to compute a single-cell quantitative phase image mainly according to the steps described in **Materials and Methods**, except a down-sampling step is performed before image reconstruction to evaluate an equivalent GVD (GVD_eq_) condition (**Materials and Methods**). The pre-generated mask (in task (1)) is used to identify the cell region.

Similar to task (2), the iTM data is first down-sampled to achieve the same GVD_eq_ condition prior to generation of the phase image (**Figure S4b, iTM**). By applying a low-pass filter on the 2D interferogram (**Figure S4c, iTM**), we obtain a bright-field contrast image; a phase-wrapped image is obtained by applying high-pass filtering (**Figure S4e(v-vi), iTM**). Finally, phase unwrapping is applied to retrieve quantitative phase (**Figure S4f, iTM**) based on the algorithm used in [2].

**S1.4 Defining GVD_eq_**

According to the time-stretch theory [1], the product of digitizer’s sampling rate and GVD critically determines the image resolution in both multi-ATOM and iTM. In iTM, extra GVD is needed to resolve the fringes in interferogram. Such condition can be represented as:

$$GVD\cdot R={2\cdot C}_{x}\cdot N_{fringe}$$

(S1)

where *R* is the sampling rate (GSa/s); GVD is the group velocity dispersion (ns/nm); C_x_ is the wavelength-to-space conversion factor (C_x_ = 6 μm/nm in our case); *N_fringe_* is the highest fringe density of interferogram (fringe/μm) according to the GVD and R. The factor of 2 in Eqn. (S1) describes the Nyquist-Shannon sampling condition, i.e. 2 samples/fringe. By this equation, we could investigate how GVD influences the fringe density and thus the image resolution in iTM.

In principle, one could physically time-stretch the optical pulses using fiber with different GVDs at a given sampling rate. However, this could introduce confounding factors complicating the analysis of GVD effect, notably the fiber loss (both coupling and dispersive loss) that influences image SNR. More importantly, such loss of SNR could result in faulty phase-unwrapping during QPI reconstruction in iTM, and thus hinders fair image comparison between multi-ATOM and iTM.

Recognizing that changing GVD at a fixed sampling rate has the equivalent effect on image resolution to varying the sampling rate at a fixed GVD [1] (Eqn. (S1)), we study the GVD effect by defining a range of equivalent GVD (GVD*_eq_*) at the fixed sampling rate *R_0_* adopted in this work (4 GSa/s), and it can be evaluated by:

$$\frac{{GVD}_{eq}}{R_{eq}}=\frac{{GVD}_{0}}{R_{0}}$$

(S2)

where *R_eq_* is the equivalent sampling rates obtained by digital down-sampling the experimental iTM and multi-ATOM image data at a fixed *GVD_0_* (= 1.78 ns/nm adopted in the current work). To ensure legitimate comparison of GVD-dependent resolution between iTM and multi-ATOM using Eqn. (S2), it is essential to first obtain simultaneously both the time-multiplexed replicas in multi-ATOM and the temporal interferogram in iTM under an oversampled condition (at 80 GSa/s). This is a critical step that ensures diffraction-limited resolution is achieved in both modalities [1] before digital down-sampling to obtain $R_{eq}$. According to the above considerations, image reconstruction and phase retrieval are applied to the down-sampled data stream at different GVD_eq_ conditions (**Figure 3a**). Based on Eqn. (S2), we could evaluate a range of GVD*_eq_* (in our case, $GVD_{eq}=0.445R_{eq}$) and investigate how GVD*_eq_* impacts the image resolution in both multi-ATOM and iTM.

**Supplementary figures**

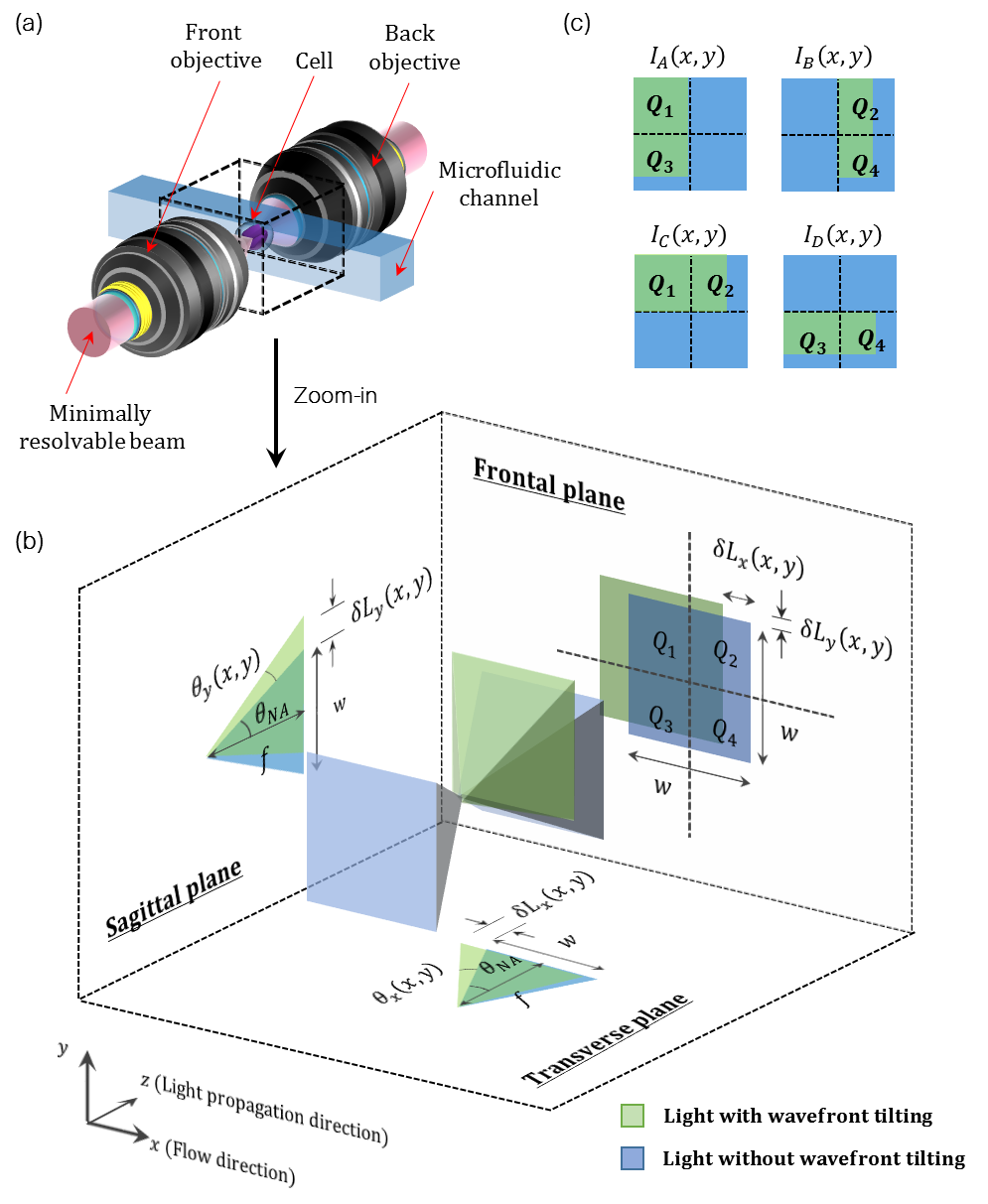

**Figure S1. Visualization of phase-gradient-induced light beam tilt.** **(a)** A minimally resolvable beam of the spectral shower is focused onto a phase object (e.g. a cell) by the front objective and collected by the back objective. **(b)** The zoom-in view of the focusing beam (assuming square beam cross-section). In the absence of phase gradient, the beam path (blue) experiences no wavefront tilt; in the presence of phase gradient, the beam path (green) is tilted. To derive the mathematical relationship between the tilt angles of wavefront ($\theta_{x}\left( x,y \right)$ and $\theta_{y}\left( x,y \right)$) and the intensity of asymmetrically-detected images ($I_{A}\left( x,y \right)$, $I_{B}\left( x,y \right)$, $I_{C}\left( x,y \right)$ and $I_{D}\left( x,y \right)$), this model considers the beam paths projected to three orthogonal planes (transverse plane, sagittal plane and frontal plane) where *x* is the cell flow direction and *z* is the light propagation direction. Transverse and sagittal planes illustrate the geometrical relationships between tilt angles of wavefront ($\theta_{x}\left( x,y \right)$ and $\theta_{y}\left( x,y \right)$) and transverse beam translation (${\delta L}_{x}$ and $\delta L_{y}$); frontal plane illustrates the intensity distribution of light at the back objective (precisely on the Fourier or far-field plane of the sample) where the area of the blue square on this plane equals to the aperture area of the back objective under the NA matching condition of two objectives. This is the plane where the knife-edge cut (i.e. asymmetric-detection) is applied. **(c)** The intensity distribution of coupled beams after four different asymmetric-detections (green portion) is illustrated ($I_{A}\left( x,y \right)$, $I_{B}\left( x,y \right)$, $I_{C}\left( x,y \right)$ and $I_{D}\left( x,y \right)$) where $Q_{1}$, $Q_{2}$, $Q_{3}$ and $Q_{4}$ refer to four quadrants of the aperture. The difference in detection portion thus gives different contrasted images of the same object. Detailed descriptions of all symbols are referred to **Materials and Methods Section 4.2**.

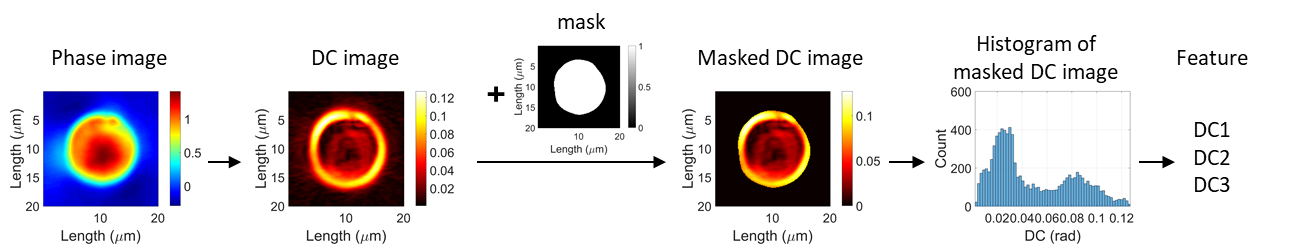

**Figure S2. Derivation of DC1, DC2 and DC3 from quantitative phase images.** Quantitative phases are transformed to dry-mass-density contrast (DC) by convolution with a kernel, within which standard deviation of phase is calculated. As a result, a DC image is transformed from a phase image. Then, a mask is applied to segment the cell body to compute the DC histogram. The first, second and third order statistical moments are calculated from the histogram which corresponds to the DC1, DC2 and DC3 respectively.

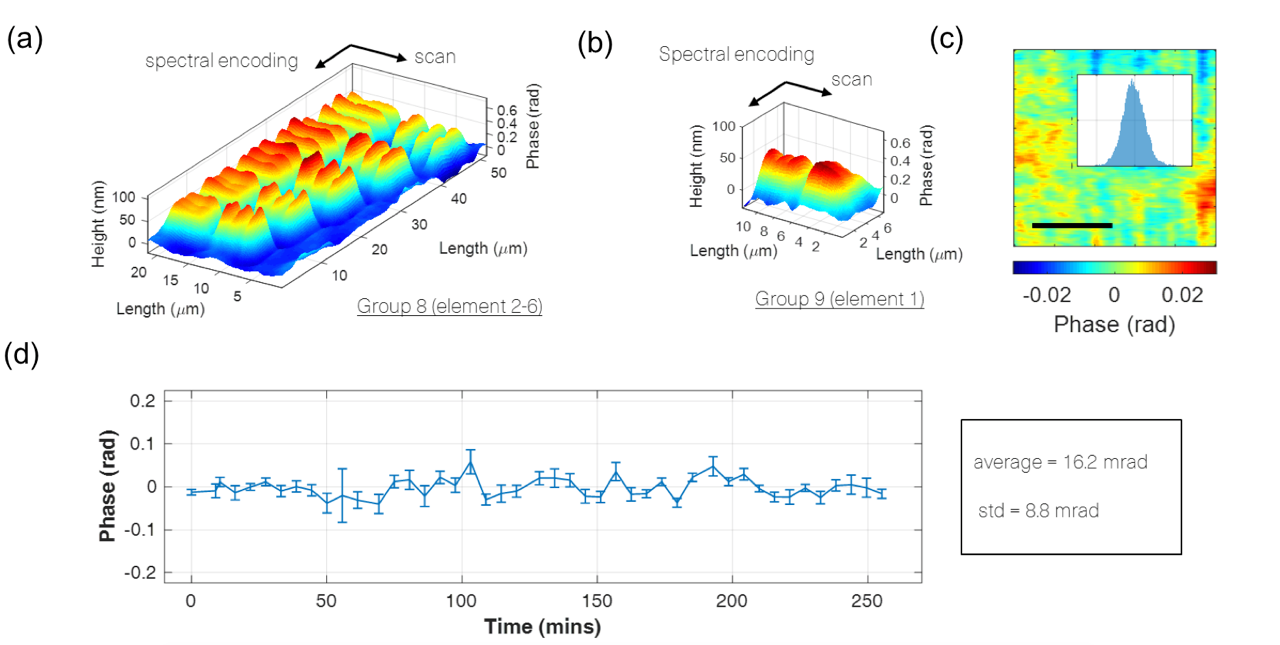

**Figure S3.** **Performance characterization. (a,b)** Quantitative phase images of the quantitative phase target with a USAF resolution chart captured by multi-ATOM (**(a)** group 8 element 2 to 6 and **(b)** group 9 element 1). **(c)** A background quantitative phase image of phosphate-buffered saline (PBS) flowing in microfluidic channel without any cells and histogram showing the corresponding phase distribution (inset). Scale bar: 10 μm. **(d)** Variation in averaged phases of quantitative phase $\phi$images of fast flowing PBS solution captured across 260 mins. std: standard deviations.

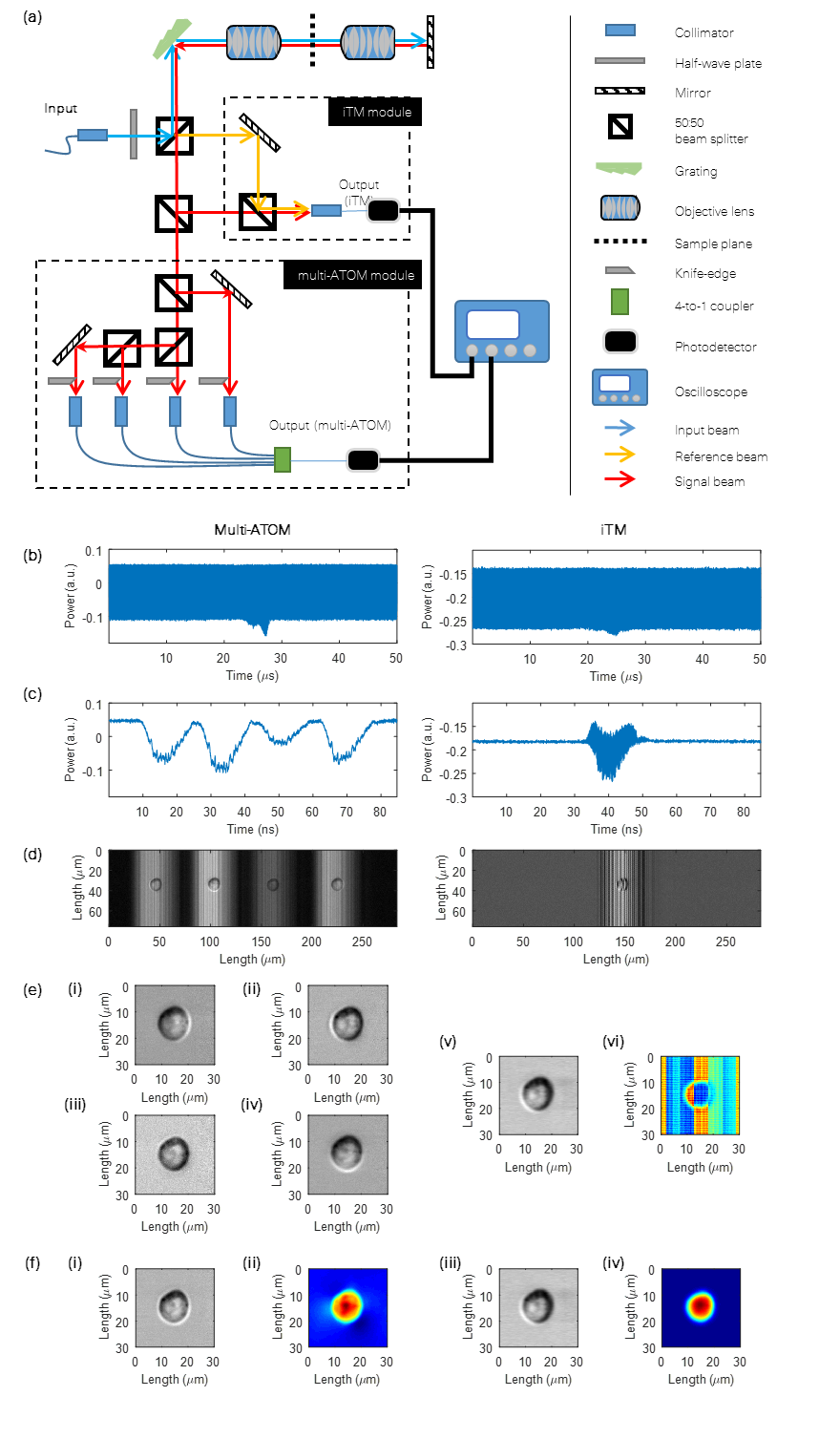

**Figure S4. System and workflow of synchronized multi-ATOM and iTM image capture.**

**Figure S4. (con’t) System and workflow of synchronized multi-ATOM and iTM image capture. (a)** Schematics of imaging system for synchronized multi-ATOM and iTM image capture. **(b)** multi-ATOM & iTM: 1D raw temporal data streams of the same cell. **(c)** Single line-scan imaging the background (multi-ATOM: four asymmetrically-detected signals; iTM: interferogram). **(d)** multi-ATOM & iTM: 2D images reconstructed by digitally stacking line scans from the raw data streams. **(e)** multi-ATOM: Four normalized DIC-like images of the same cell are simultaneously reconstructed (i-iv). iTM: Bright-field (v) and wrapped-phase (vi) are obtained by applying low-pass and high-pass filtering respectively. **(f)** Digitally reconstructed images: (i, iii) bright-field and (ii, iv) quantitative phase images. See also **Supplementary Methods S1.3.**

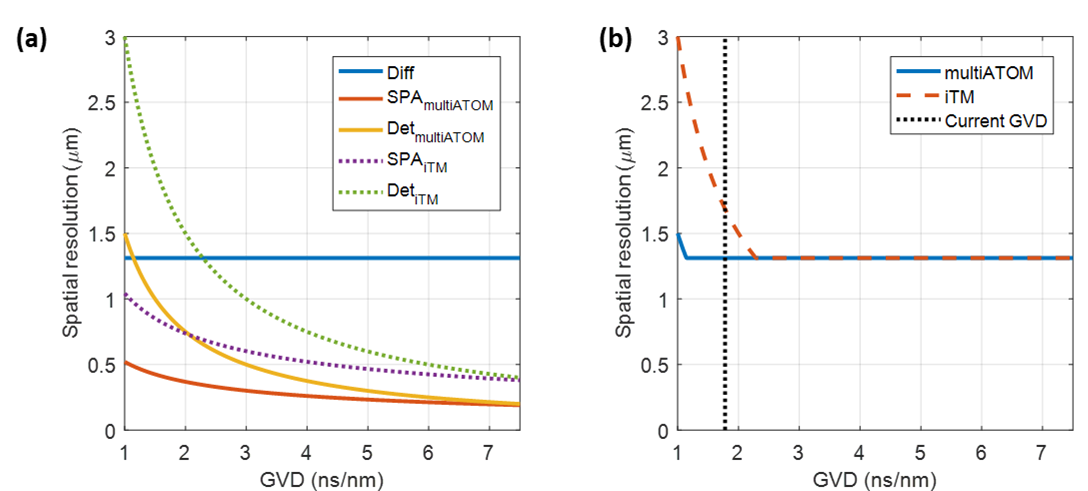

**Figure S5.** **Time-stretch image resolution analysis of multi-ATOM and iTM. (a)** Overview of relationships between group-velocity-dispersion (GVD) and three resolution constrains of time-stretch microscopy (i.e. diffraction-limited (Diff), SPA-limited (SPA) and detector-limited (Det)). Due to an extra modulation by interferogram, iTM requires at least a double of sample point to resolve the same object, thus, showing a reduction in SPA-limited and detector-limited resolutions by half. **(b)** The best achievable resolution of both multi-ATOM and iTM at different GVD summarized from (a) are shown. Under the current GVD (= 1.78 ns/nm, black dotted line) and sampling rate (= 4 GHz), only multi-ATOM can achieve diffraction-limited resolution (~1.31 μm) while iTM can only achieve detector-limited resolution (~1.69 μm), except increase the GVD at least to 2.29 ns/nm that is not commercially available.

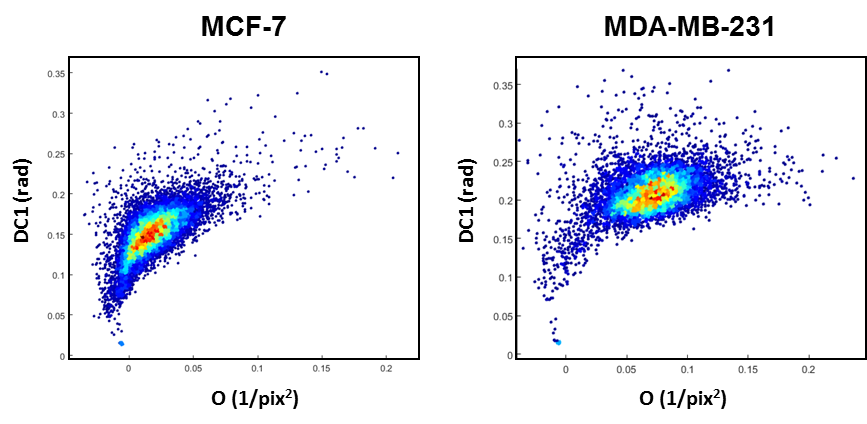

**Figure S6. Pure populations of MCF-7 and MDA-MB-231 measured by multi-ATOM.** Bivarate plots based on two features derived from multi-ATOM images of MCF-7 and MDA-MB-231. Two pure cell populations were imaged separately.

**Supplementary Table**

| Name | Symbol | Unit | Equation |
| --- | --- | --- | --- |
| Normalized light intensity | I | - | - |
| Quantitative phase | $\phi$ | rad | - |
| Spatial coordinate | *x, y, a, b* | - | - |
| Length of binary mask | $l$ | μm | - |
| Width of binary mask | w | μm | - |
| Number of pixel in binary mask | $n_{m}$ | **-** | ***­*-** |
| Pixel-to-length conversion factor | $C_{p\to s}$ | μm | - |
| Center wavelength of light | $\lambda$ | nm | $1064$ |
| Specific refractive increment | $\alpha$ | ml/g | $0.2$ |
| Perimeter | p | μm | $\int_{0}^{2\pi} \sqrt{\left( \left( \frac{dx}{dt} \right)^{2}+\left( \frac{dy}{dt} \right)^{2} \right)} dt$ |
| Area | A | μm^2^ | $C_{p\to s}\cdot n_{m}$ |
| Volume | Vol | μm^3^ | $\left( \pi\cdot w^{2}\cdot l \right)/6$ |
| Roundness | R | - | $4\pi\cdot A/p$ |
| Opacity | O | 1/ μm^2^ | ${\iint_{A} \left( 1-I\left( x,y \right) \right) dxdy}/A$ |
| Mean of normalized light intensity | $\bar{I}$ | - | ${\iint_{A} I\left( x,y \right) dxdy}/{n_{m}}$ |
| Intensity variance | $\sigma_{I}^{2}$ | - | ${\iint_{A} \left( I(x,y)-\bar{I} \right)^{2} dxdy}/\left( n_{m}-1 \right)$ |
| Intensity skewness | $\gamma_{I}$ | - | $\frac{{{\iint_{A} \left( I\left( x,y \right)-\bar{I} \right)^{3} dxdy}/n}_{m}}{\left( \sqrt{{\iint_{A} \left( I\left( x,y \right)-\bar{I} \right)^{2} dxdy}/{n_{m}}} \right)^{3}}$ |
| Maximum phase | $\phi_{max}$ | rad | $\max\left\{ \phi\left( x,y \right) \right\}$ |
| Mean of phase | $\bar{\phi}$ | rad | ${\iint_{A} \phi\left( x,y \right) dxdy}/{n_{m}}$ |
| Phase variance | $\sigma_{\phi}^{2}$ | rad^2^ | ${\iint_{A} \left( \phi(x,y)-\bar{\phi} \right)^{2} dxdy}/{(n_{m}-1)}$ |
| Phase skewness | $\gamma_{\phi}$ | rad | $\frac{{{\iint_{A} \left( \phi\left( x,y \right)-\bar{\phi} \right)^{3} dxdy}/n}_{m}}{\left( \sqrt{{\iint_{A} \left( \phi\left( x,y \right)-\bar{\phi} \right)^{2} dxdy}/{n_{m}}} \right)^{3}}$ |

**Table S1. Definitions of symbols and equations of the biophysical features extracted from multi-ATOM.**

| Name | Symbol | Unit | Equation |
| --- | --- | --- | --- |
| Dry mass | DM | pg | $\frac{\lambda}{2\pi\cdot\alpha}\iint_{A} \phi(x,y) dxdy$ |
| Dry mass density | DMD | pg/fl | $DM/Vol$ |
| Kernel width | r | - | - |
| Number of pixel in kernel | $n_{k}$ | - | $r^{2}$ |
| Mean of phase within kernel | $\bar{\phi}_{k}(x,y)$ | rad | ${\int_{y-\frac{r}{2}}^{y+\frac{r}{2}} \int_{x-\frac{r}{2}}^{x+\frac{r}{2}} \phi\left( a,b \right) dadb}/{n_{k}}$ |
| Local standard deviation of phase | $\phi_{L}$/DC | rad | $\sqrt{{\int_{y-\frac{r}{2}}^{y+\frac{r}{2}} \int_{x-\frac{r}{2}}^{x+\frac{r}{2}} \left( \phi\left( a,b \right)-\bar{\phi}_{k}\left( x,y \right) \right)dadb}/\left( n_{k}-1 \right)}$ |
| Dry-mass-density contrast 1 | DC1/$\bar{\phi}_{L}$ | rad | ${\iint_{A} \phi_{L}\left( x,y \right) dxdy}/{n_{m}}$ |
| Dry-mass-density contrast 2 | DC2 | rad | $\sqrt{{\iint_{A} \left( \phi_{L}\left( x,y \right)-\bar{\phi}_{L} \right)^{2}dxdy}/\left( n_{m}-1 \right)}$ |
| Dry-mass-density contrast 3 | DC3 | rad | $\frac{{\iint_{A} \left( \phi_{L}(x,y)-\bar{\phi}_{L} \right)^{3} dxdy}/{n_{m}}}{\left( \sqrt{{\iint_{A} \left( \phi_{L}\left( x,y \right)-\bar{\phi}_{L} \right)^{2} dxdy}/{n_{m}}} \right)^{3}}$ |

**Table S1 (con’t). Definitions of symbols and equations of the biophysical features extracted from multi-ATOM.**
